# Midbrain dopamine neurons encode internal effort signal

**DOI:** 10.1101/2020.03.04.976969

**Authors:** Dong V Wang

## Abstract

Midbrain dopamine neurons of the ventral tegmental area (VTA) play a central role in both positive and negative motivation. As motivation and effort are intrinsically connected, we asked how the VTA dopamine neurons may also process internally driven effort signal. We designed two novel jumping tasks, the reward- and escape-motivated jumps, and revealed that the VTA dopamine neurons exhibit an effort-correlated activation during both jumping behaviors.

## Results

Growing evidence suggests that dopamine is essential for regulating effort-related brain functions and behaviors^1-3^. However, how the midbrain dopamine neurons may encode such effort signal remains unclear. Most research to date has been confined to study the dopamine neuron’s responses to external stimuli, either rewarding^4^ or aversive^5-8^. Fewer studies have examined how the dopamine neurons react during internally guided behaviors^9-14^. Here we asked how the ventral tegmental area (VTA) dopamine neurons may process internally driven effort signal.

We implanted one bundle of 8 tetrodes (32 channels) into the VTA of each mouse, and the recording sites were later confirmed by histology (Figure 1A). A total of 11 VTA putative dopamine neurons were classified (see Methods) and thus used for analysis. These putative dopamine neurons exhibited relatively low baseline firing rates (5.2 ± 1.2 Hz, mean ± SD) and significant activations in response to a reward-predicting tone (Figure 1 A&D, bottom). In addition, all tested putative dopamine neurons were confirmed to be suppressed (<20% baseline, see Methods; Figure 1D, top) by apomorphine, an agonist that inhibits dopamine neuron activity through its autoreceptors^15^.

**Figure 1.**
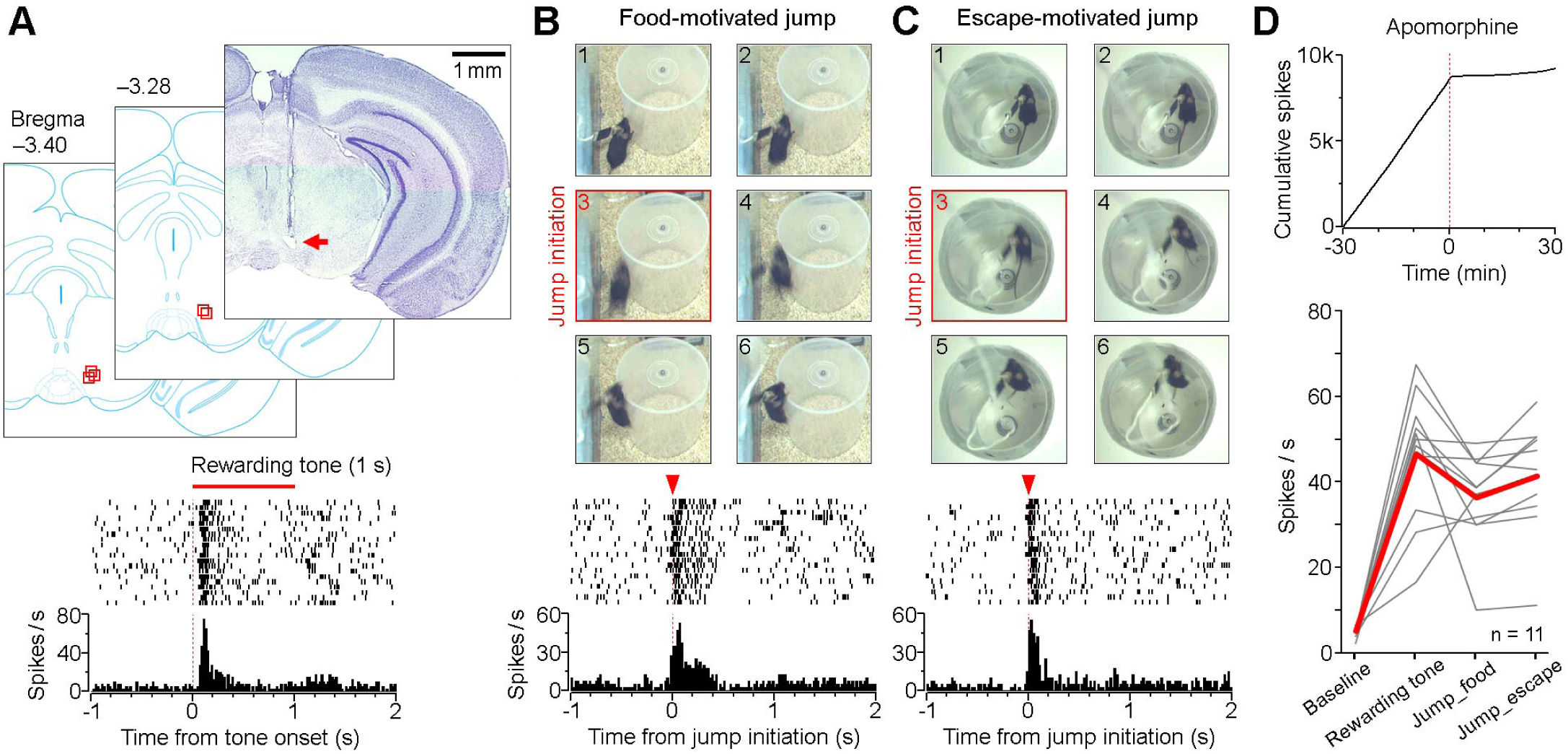
Convergent processing of reward and effort signals by the VTA dopamine neuron. (A) Top, a representative coronal section and schematics showing the recording sites in the VTA (arrow and red squares). Bottom, peri-event rasters (20 trials) and histogram of a putative dopamine neuron in response to the rewarding tone that predicted sugar pellets (delivered at the end of each tone). (B&C) Top, six consecutive frames (33 ms per frame) showing reward-motivated (B) and escape-motivated (C) jumping behaviors. Frames #3 were defined as the jump initiation based on high motion (see Methods). Bottom, peri-event rasters (20 trials) and histograms of the same neuron (as shown in A) during reward- (B) and escape-motivated (C) jumping behaviors. (D) Top, cumulative spikes of the same neuron (as shown in A) in response to apomorphine (1 mg/kg; i.p. injection at time 0). Bottom, baseline and peak firing rates of individual putative dopamine neurons (grey lines; red line indicates the mean) at three conditions, that is, in response to the rewarding tone, at the initiation of food- and escape-motivated jumps. There’s no difference of the peak firing rate among the three conditions (P = 0.20, repeated measures ANOVA).

To determine how the dopamine neurons may encode effort signal, we designed two high-effort jumping tasks, the reward- and escape-motivated jumps. During the tasks, animals either voluntarily jumped on a high platform to receive sugar pellets (Figure 1B) or jumped to escape an aversive chamber that infrequently quaked^8^ (Figure 1C; see Methods). This jump initiation was completely voluntary with no immediate external trigger; therefore, the coincided neural activity likely represented an internal effort signal associated with jump initiation. Our results revealed that the VTA putative dopamine neurons exhibited robust burst firing at the initiation of both jumping behaviors (Figure 1 B–D, bottom; Suppl. figure 1). This corroborates with recent studies that reported an activation of substantia nigra pars compacta (SNc) dopamine neurons at movement initiation^11,12^. However, unlike the SNc where distinct dopamine populations respond to food reward versus movement initiation effort^11,12^, the same VTA dopamine population processes both reward and jumping effort information convergently.

We next sought to compare dopamine activity during successful versus failed jumps, termed higher- and lower-effort jumps, respectively. A success was defined if the animal initiated a jump and reached its goal (either food or escape); otherwise a failed attempt was defined as a failure (see illustrations in Figure 2). We rationalized that VTA dopamine neurons increase firing whenever an animal makes an effort to jump. We also expected to see a higher dopamine activation in successful than failed jumps. In agree with these predictions, our results revealed that VTA putative dopamine neurons exhibited an activation at the initiation of all jumps, with a significant higher activity in successful compared to failed trials (Figure 2 A–D). This gradient activation indicates an encoding of effort magnitudes by the VTA dopamine neurons.

**Figure 2.**
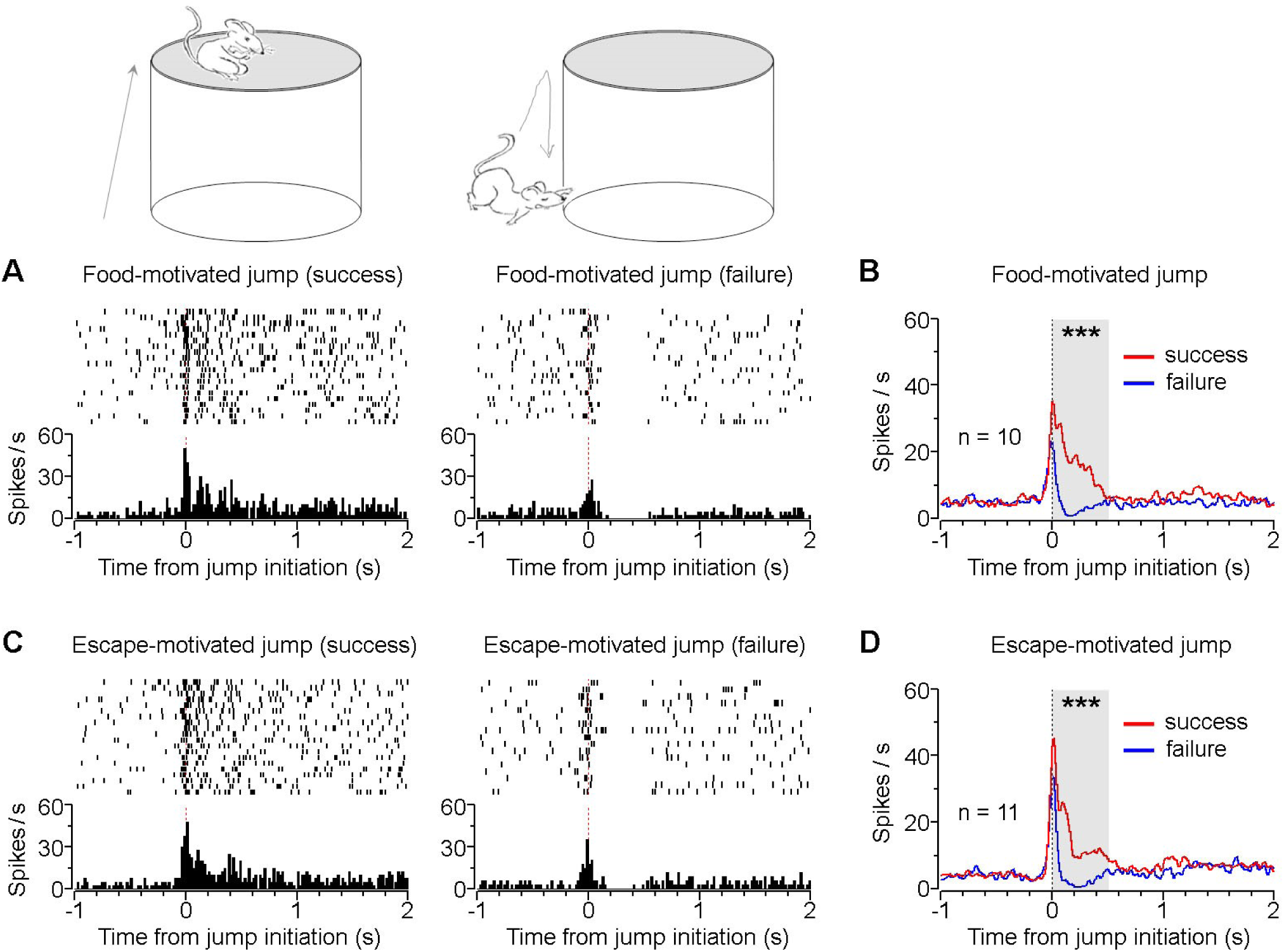
Successful and failed jumps. (A) peri-event rasters and histogram of a VTA putative dopamine neuron during successful (left) or failed (right) reward-motivated jumps. (B) Mean activity of VTA putative dopamine neurons (n = 10) during successful and failed reward-motivated jumps. (C) peri-event rasters and histogram of the same neuron (as shown in A) during successful (left) or failed (right) escape-motivated jumps. (D) Mean activity of VTA putative dopamine neurons (n = 11) during successful and failed escape-motivated jumps. Only data from the 15-cm jumps were used for analysis here (see Figure 3). ***P < 0.001, unpaired t-test of the means between 0–0.5 s.

To more directly test how the dopamine neurons process jumping effort magnitudes, we gradually elevated the platform/chamber height (13→15→17 cm; see Methods). Behaviorally, the mice showed high success rate when the jumping height was 13 cm (71% and 74% for reward- and escape-motivated jumps, respectively). However, the success rate dropped markedly when the height was slightly elevated to 15 cm (46%; 56%) or 17 cm (14%; 25%). Given the low success rate at 17 cm, we combined the data from the 15- and 17-cm jumps for analysis. Our results revealed that VTA putative dopamine neurons exhibited more sustained activity during the higher-compared to lower-effort jumps (Figure 3 A–D). The lasting activity between ∼0.1–0.5 s likely corresponded to the climbing effort that immediately followed the initial jumping effort (between 0–0.1 s; see Methods).

**Figure 3.**
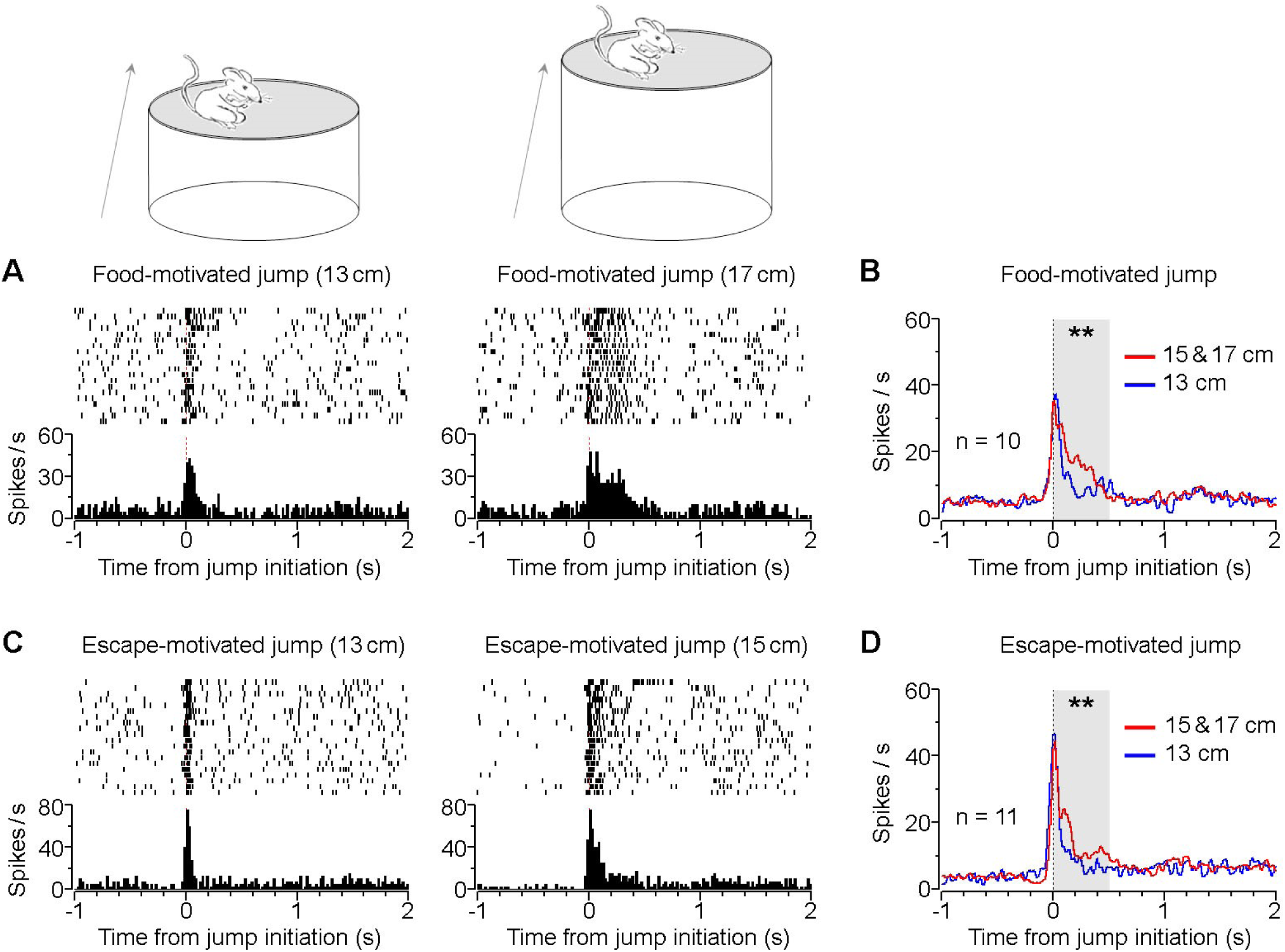
VTA dopamine neurons exhibit effort-correlated activation. (A) peri-event rasters and histogram of a VTA putative dopamine neuron during reward-motivated jumps at a height of 13 cm (left) and 17 cm (right). (B) Mean activity of VTA putative dopamine neurons (n = 10) during reward-motivated jumps at a higher (red) and lower height (blue). (C) peri-event rasters and histogram of the same neuron (as shown in A) during escape-motivated jumps at a height of 13 cm (left) and 15 cm (right). (D) Mean activity of VTA putative dopamine neurons (n = 11) during escape-motivated jumps at a higher (red) and lower height (blue). **P < 0.01, unpaired t-test of the means between 0–0.5 s.

Animals’ actions and behaviors are rarely totally externally driven, nor are ever completely internally guided. Rather, they often comprise both external and internal components. In our experiments, animal’s jump behavior was voluntary and internally driven by positive anticipation, but not triggered by any immediate external stimuli. Therefore, we suggest that the VTA dopamine activity at jump initiation represented, to a greater extent, an internal effort signal that was associated with anticipated positive outcomes. Importantly, this dopamine activity correlated well with effort magnitude: the higher effort the jumps, the higher activation the dopamine neurons. Together with recent studies that report dopamine activation at movement initiation^9-13^, we suggest that midbrain dopamine neurons are essential in processing an internal effort signal that influences relevant downstream cortical/subcortical regions in invigorating goal-directed behaviors^16^.

## Methods

### Subjects

Male mice (C57BL/6J; Jackson Lab) were individually housed in customized homecages (40 × 20 × 25 cm) on a 12-h light/dark cycle after electrode implant surgery (∼3 months old at the time of surgery). Spike data from five mice were used for analysis. All procedures were approved by the Institutional Animal Care and Use Committee, Medical College of Georgia, Augusta University.

### Surgeries

We constructed a 32-channel (8 tetrodes), ultra-light (weight ∼1 g), adjustable (screw-driven) electrode bundle, similar to that described previously^8,9^. Each tetrode consisted of four 18-µm diameter wires (90% Platinum 10% Iridium, California Fine Wire). On the day of surgery, mice were anesthetized with Ketamine/Xylazine (80/12 mg/kg, i.p.); the electrode bundle was then lowered towards the VTA in the right hemisphere (AP –3.4 mm; ML 0.5 mm; DV ∼3.9 mm) and secured with dental cement.

### Tetrode recording

About three days after surgery, electrodes were screened daily for neural activity. If no putative dopamine neuron was detected, the electrode bundle was advanced ∼50 µm daily, until we recorded one putative dopamine neuron^8^. Spikes (filtered at 250–8k Hz; digitized at 40 kHz) were recorded during the entire experimental process using the Plexon acquisition system (Plexon Inc.). Mice behaviors were simultaneously recorded using Plexon CinePlex (30 frames/s). Recorded spikes were sorted using Plexon OfflineSorter^8^.

### Reward conditioning

Mice were slightly food restricted and habituated to sugar pellets a few days before training. During conditioning, mice were trained to pair a tone (5 kHz, 1 s; ∼80 dB) with subsequent sugar pellet delivery for at least two days (40–60 trials per day; 1–2 min interval between trials). The tone was generated by the A12-33 audio signal generator (5-ms shaped rise/fall; Coulbourn Instruments). A sugar pellet (14 mg; Bio-Serv) was delivered by a food dispenser (Med. Associates Inc.) and dropped into a food receptacle at the termination of the tone^8^.

### Reward- and escape-motivated jumps

All mice were previously trained with the reward conditioning (see above) and slightly food restricted. To induce reward-motivated jump, we first placed a platform (10 cm in diameter; ∼10 cm in height) in the homecage, and meanwhile, we placed a sugar pellet (14 mg) at the platform center. Mice often grasped the edge of the platform and climbed on to get the sugar pellet. After consumption of the pellet, the mouse either voluntarily got down of the platform or was guided down by the experimenter. Then another food pellet was placed on the platform to trigger the next climbing trial. After 15–20 trials, we gradually elevated the platform height to 13, 15 and 17 cm. Typically, the mice were able to climb on the platform when the height was 12 cm or lower. However, when the height exceeded 12 cm, the mice had to jump to reach the edge of the platform. This initial jump (lasting ∼0.1 s) was followed by an effort-based climbing onto the platform (lasting up to 0.5 s; see Figure below). Images were visually screened frame by frame (33 ms per frame), and the jump initiation was defined as the first blurry frame (due to the high motion of jumps; see Figure below).

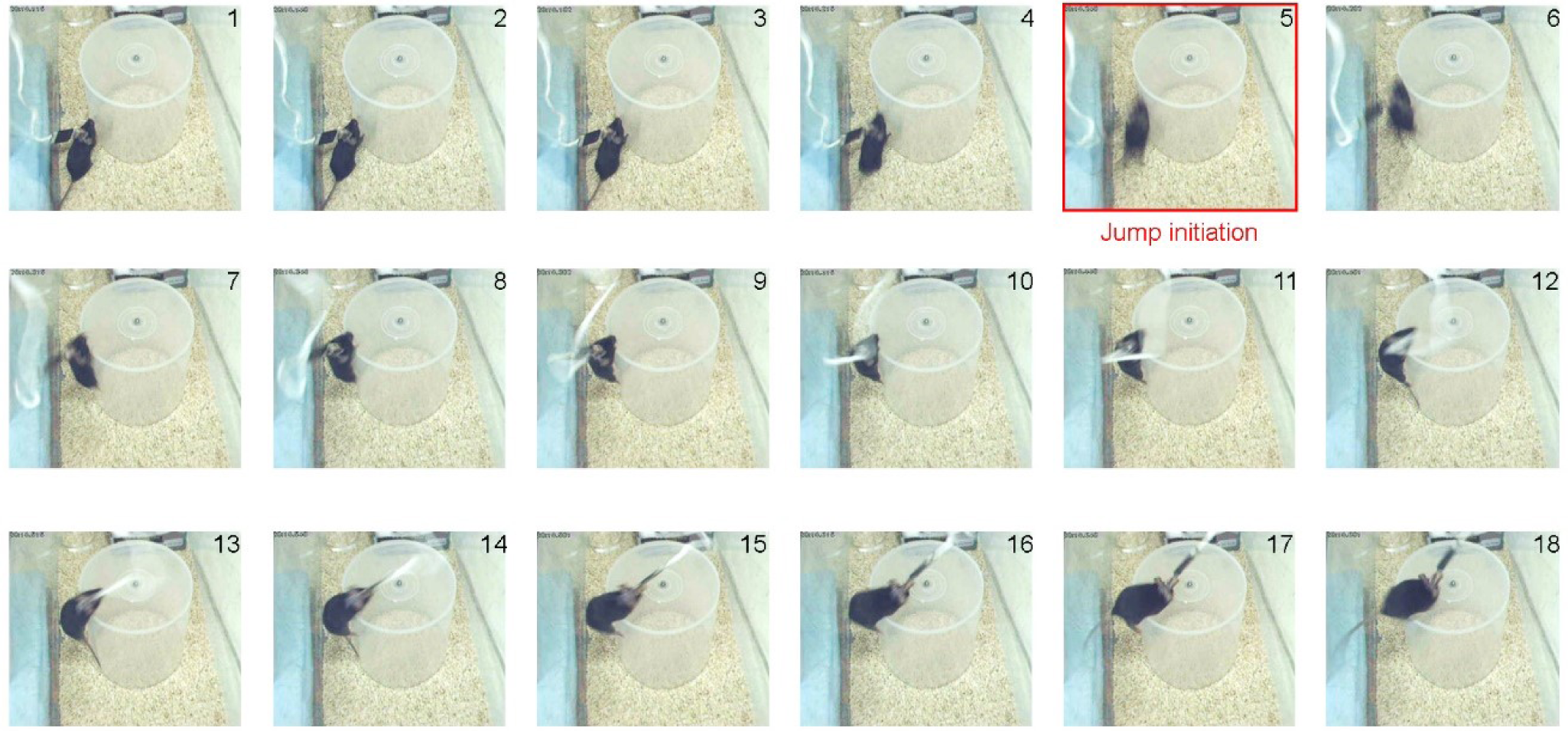

To induce escape-motivated jump, we first placed the mouse into a small chamber (10 cm in diameter; 13 cm in height) and infrequently shook the chamber (1–2 min interval between trials). All putative dopamine neurons showed suppressed activity during the shake events, corresponding to the Type-I/II neurons described previously^8^. After about 5–10 minutes, the mice typically initiated jumps to escape the chamber. Mice were manually placed back to the chamber to trigger the next trial of jump. Similarly, we gradually elevated the height to 15 and 17 cm after 15–20 trials at each height.

### Data Analysis

Dopamine neurons were classified based on previously established criteria^8^. Briefly, all classified dopamine neurons (n = 11) exhibited a low baseline firing rate (2.3–6.5 Hz) and long inter-spike interval (>4 ms). In addition, 10 of the 11 neurons showed significant activations in response to the reward-predicting tone (Figure 1D; P < 0.001; Wilcoxon signed-rank test)^8^. Six of them were tested with apomorphine (1 mg/kg, i.p.) and all six showed a robust suppression (<20% baseline during the first half an hour). Peri-event rasters and histograms were conducted in NeuroExplorer (Nex Technologies).

### Histology

On completion of the experiments, the final electrode position was marked by passing a 10-s, 20-µA current (Stimulus Isolator A365, WPI) through two electrodes. Mice were deep anesthetized and perfused with 0.9% saline followed by 4% paraformaldehyde (PFA). Brains were then removed and post-fixed in PFA for 24 h or longer. Brains were rapidly frozen and sliced on a cryostat (50 µm) and stained with cresyl violet.

## Acknowledgements

I thank Dr. Joe Z Tsien for the support of this project and Kun Xie for assistance with the microdrive construction. Data used in this paper was collected by Dong V Wang under the supervision of Joe Z Tsien at the Medical College of Georgia, Augusta University between 2008–2011. I also thank Dr. Satoshi Ikemoto for comment on an earlier draft.

---

**Suppl. figure 1.**
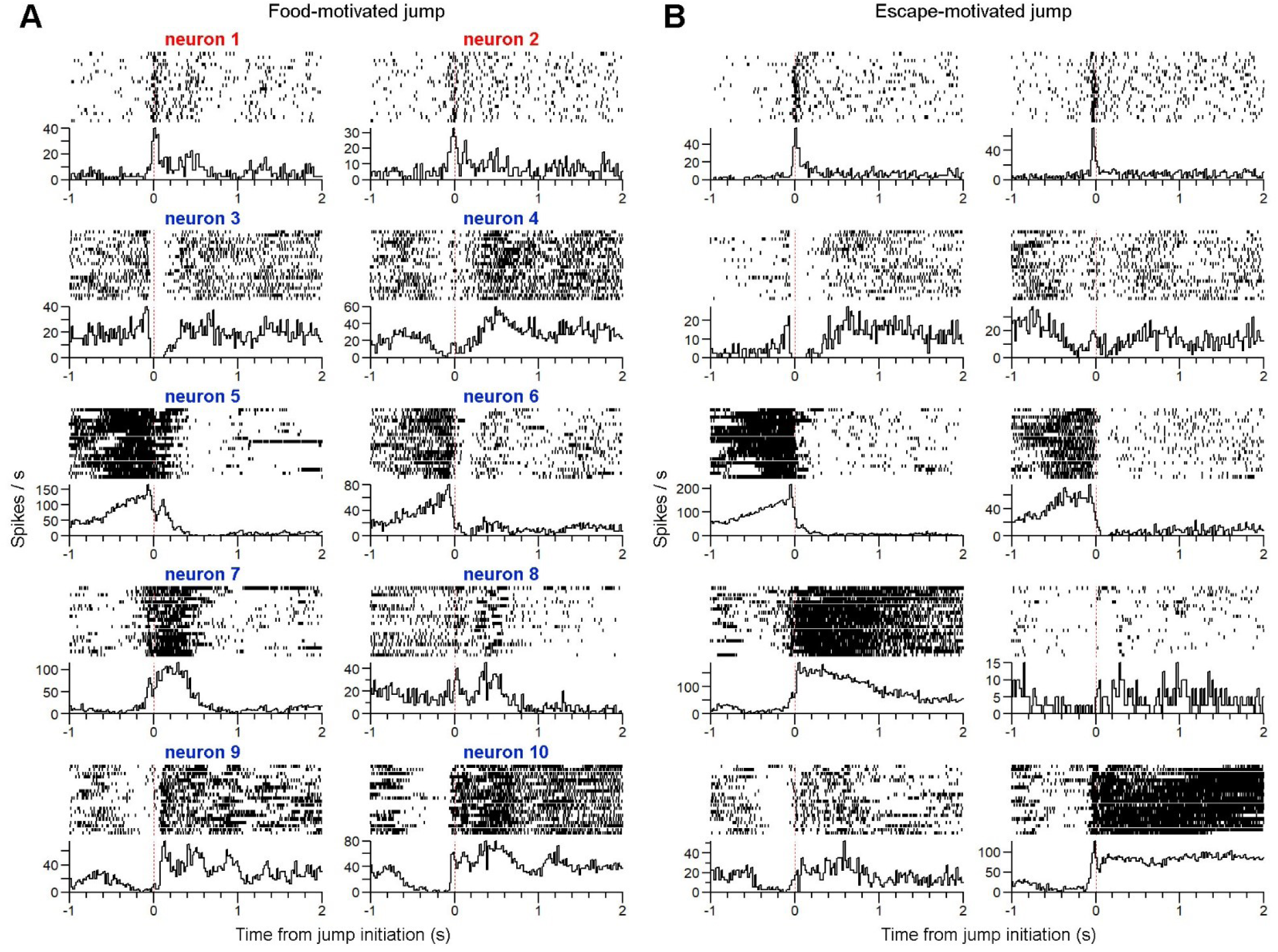
Activity of VTA putative dopamine and non-dopamine neurons during jump behaviors. (A&B) peri-event rasters and histograms of 10 simultaneously recorded VTA neurons during food- (A) and escape-motivated (B) jumps. Neurons 1 & 2 are putative dopamine neurons, whereas neurons 3–10 are putative non-dopamine neurons.

